# Exact Expectation of Complete Spatial Randomness for Nearest Neighbor *G*(*r*): A Scalable Alternative to Permutations

**DOI:** 10.1101/2025.06.11.659088

**Authors:** Alex C Soupir, Brandon J Manley, Lauren C Peres, Brooke L Fridley, Julia Wrobel

## Abstract

Spatial analysis is becoming increasingly important for studies, from epidemiology to tissue biology, as technologies advance and experimental costs decrease. However, the widespread use of spatial metrics such as Nearest Neighbor *G*(*r*) is affected by the fact that biological systems rarely satisfy the assumption of stationarity, which is required to appropriately use theoretical complete spatial randomness (CSR) measures. As a result researchers often use computationally expensive permutations to empirically estimate CSR for subsets of points or cells. Here, we present closed form analytical solutions for both the mean and variance of the sample-specific CSR for Nearest Neighbor *G*(*r*) to allow for fast and reproducible calculation without permutations. Using a multiplex immunofluorescence sample of clear cell renal cell carcinoma, we show that the theoretical *G*(*r*) for cytotoxic T cells overestimates CSR at low radii (due to spatial constraints between cells) while drastically underestimating CSR at radii between 20 and 90 pixels. In a simulated sample of 30 points, our analytical solution for the mean is identical to the average of *G*(*r*) measured on all 142,506 unique combinations of 5 marked (or positive) points. On the real clear cell renal cell carcinoma sample, our exact CSR is similar in speed to estimating CSR with 1000 permutations while our optimized Rcpp implementation is ∼30x faster and consuming ∼20x less memory than 1000 permutations. This permutation-free approach dramatically enhances computational efficiency and reproducibility, enabling scalable and reproducible analysis for studies in epidemiology, multiplex immunofluorescence, spatial transcriptomics, and related fields where accurate, sample-specific null expectations are important for comparisons.

## Introduction

The analysis of spatial point patterns is fundamental across multiple disciplines, including ecology, epidemiology, and spatial omics technologies such as multiplex immunofluorescence and spatial transcriptomics [1–7]. Understanding how objects are distributed in space can provide critical insights into biological organization, environmental interactions, and population dynamics. One widely used method for assessing the spatial distribution of observed points is the Nearest Neighbor *G*(*r*) function, which provides insight into clustering and dispersion by calculating the cumulative distribution of nearest-neighbor distances [1,2,6,8–12]. By measuring the likelihood that a point’s nearest neighbor falls within a given distance, the function helps distinguish random, clustered, or overdispersed spatial arrangements. This metric is routinely applied in diverse domains such as assessing the spatial organization of immune cells in tumors, studying plant competition in ecological systems, and mapping the spread of infectious diseases. However, the widespread application of the Nearest Neighbor *G*(*r*) function faces a critical limitation: the assumption that the underlying point pattern follows complete spatial randomness (CSR) is frequently violated in real-world datasets [6]. Many biological systems exhibit spatial heterogeneity (i.e., regions between tissues, distribution of proteins in cells, elements such as bodies of water in mapping or blood vessels in tissue) making traditional CSR-based expectations unsuitable for comparison. Importantly, studies such as those from epidemiology and spatial omics analysis have multiple samples which makes it critical to adjust for sample-specific effects such as deviations from complete spatial randomness when comparing across samples.

To address this issue, researchers typically employ a permutation-based approach to estimate a sample-specific null distribution, which accounts for spatial heterogeneity by randomly reassigning observed marks (e.g., cell types, species) to spatial locations and recalculating *G*(*r*) for each permutation [6]. By averaging across many permutations, an empirical estimate of the expected function under CSR is obtained, which can then be used to adjust the observed *G*(*r*). While effective, this approach carries a significant computational cost, as the number of required permutations increases with sample size, data complexity, and the degree of deviation from CSR. The necessity of running hundreds or even thousands of permutations to obtain stable estimates presents a major barrier in studies that involve large-scale datasets, high-resolution spatial profiling, or iterative analysis across multiple experimental conditions. As datasets in spatial omics and epidemiology continue to grow in both volume and complexity, the limitations of permutation-based methods will become increasingly prohibitive. The reliance on repeated resampling not only demands substantial computational resources but also introduces variability in estimation, as results depend on the number of chosen permutations.

In this work, we introduce a novel approach that eliminates the need for explicit permutation enumeration by deriving a direct analytical solution for the expected *G*(*r*) function under CSR. Previously, Wrobel and Song (2024) proposed a solution for permutation-free RIpley’s K [13] By leveraging properties of the spatial point process, our method allows for the efficient estimation of the null distribution without computationally intensive simulations. This approach provides a mathematically exact expectation, reducing the dependence on iterative resampling while improving computational efficiency. The ability to compute an exact CSR expectation offers several key advantages, including improved accuracy, enhanced reproducibility, and scalability to large datasets. By bypassing the need for randomization, our method ensures that CSR estimates remain consistent across studies, removing the variability introduced by finite permutation counts. Furthermore, the computational gains from eliminating permutations make high-throughput spatial analyses feasible, allowing researchers to incorporate CSR-adjusted clustering assessments into large-scale studies without prohibitive processing times.

Our approach is broadly applicable across domains where point pattern analysis is used and points of interest are a subset of all points measured, including but not limited to epidemiology, cell biology, and cancer research; fields that are interested in assessing Nearest Neighbor *G*(*r*) measures between samples. By removing the reliance on permutation-based estimation, this method facilitates efficient spatial analyses, enhances computational reproducibility (reduced variation due to randomness in permutations), and enables the study of complex spatial phenomena with increased productivity. We demonstrate the effectiveness of this approach using spatial proteomic and transcriptomic data, but the framework extends naturally to any field in which spatial point patterns and their interactions are of interest. Given the growing importance of spatially resolved datasets in scientific research, ensuring that clustering assessments remain both statistically rigorous and computationally feasible is critical. The exact CSR method proposed here meets this need by providing an analytically derived, scalable, and efficient alternative to permutation-based CSR estimation.

## Solution Explanation

The Nearest Neighbor G-function, *G*(*r*), is a fundamental spatial statistic used to quantify clustering by evaluating the proportion of points whose nearest neighbor lies within a given radius *r*. Traditionally, an observed *G*(*r*) function is compared to a null model under CSR, which is often estimated via computationally intensive permutations. Here, we present an exact derivation for the expected *G*(*r*) under CSR, eliminating the need for permutations and significantly reducing computational overhead.

### 1. Defining the Nearest Neighbor Function

Consider a dataset consisting of *N* total points in a metric space (e.g., a 2D plane). A subset *S* of size *n* (e.g., 150 marked points) is drawn uniformly at random from these *N* points. The nearest neighbor function for a specific subset *S* is defined as:

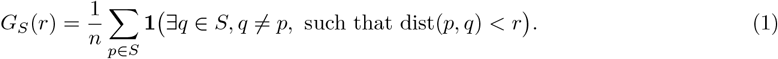

That is, for each point *p* in the subset *S*, the indicator function evaluates whether there exists another point *q* in *S* within radius *r*, and the function averages this over all *n* points in *S*.

### 2. Solution for Mean

#### 2.1 Goal

The goal is to compute the expected value of *G*_*S*_(*r*), denoted as:

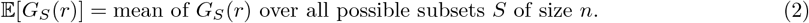

Since the number of such subsets is astronomically large (i.e.,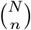), direct enumeration is infeasible. However, leveraging linearity of expectation, we derive an exact formula that can be evaluated efficiently.

#### 2.2. Reformulating Expectation Using Indicator Functions

By expanding the summation inside the expectation:

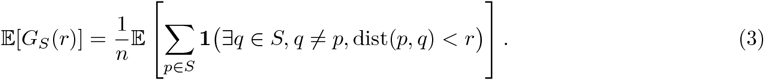

Rewriting the summation over all points *p* in the full dataset *X*, we obtain:

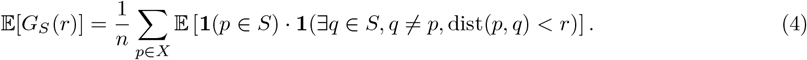

Applying linearity of expectation, this simplifies to:

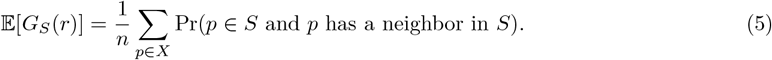

Thus, the problem reduces to computing the probability that an arbitrary point *p* is included in *S* and has at least one neighbor in *S*.

#### 2.3 Computing the Probability of Inclusion and Neighbor Presence

##### 2.3.1 Probability that *p* is in *S* Since each subset *S* of size *n* is chosen uniformly at random

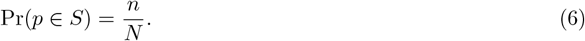

##### 2.3.2 Probability that *p* has at least one neighbor in *S*, given *p ∈S* Define *n*_*p*_ as the number of other points in *X* that lie within distance *r* of *p*

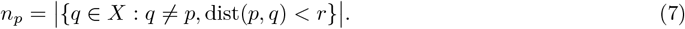

Given that *p ∈S*, the remaining *n −*1 points in *S* are chosen from *X \ {p}*, which consists of *N −*1 total points. The probability that none of the *n*_*p*_ neighbors of *p* are selected into *S* is:

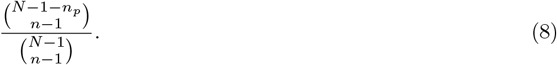

Thus, the probability that at least one of the *n*_*p*_ neighbors is included in *S* is:

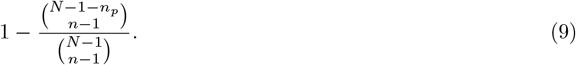

##### 2.3.3 Final Probability Computation

Since the events of selecting *p* and selecting a neighbor are independent,

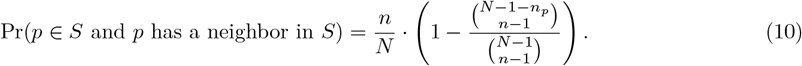

#### 2.4. Summation Over All Points

Summing over all points *p ∈ X*, we obtain:

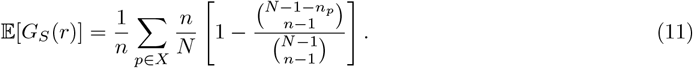

Noting that 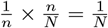, the final result simplifies to:

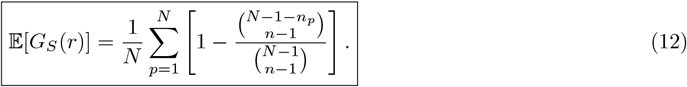

#### 2.5. Computational Considerations

- The term *n*_*p*_ (counting neighbors within *r*) must be precomputed for each point *p*, which is typically *O*(*N* ^2^) in brute force but can be optimized to *O*(*N* log *N*) with spatial data structures.
- Computing the binomial coefficient ratio directly can lead to numerical instability. A practical implementation should use log-factorial methods to evaluate:

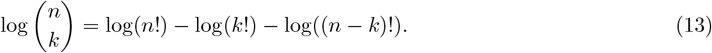

This avoids overflow when handling large factorials.

This derivation provides an exact formula for the expected *G*(*r*) function under CSR, eliminating the need for computationally expensive permutations. The final equation allows efficient computation of the CSR expectation for large-scale spatial datasets, facilitating accurate and reproducible spatial clustering analyses.

In the next section, we provide a closed-form solution for variance.

### 3. Solution for Variance of *G*_*S*_(*r*)

The variance of *G*_*S*_(*r*), the nearest-neighbor function for a random subset *S* of size *n*, quantifies the uncertainty in spatial clustering estimates under CSR. While the expectation of *G*_*S*_(*r*) provides an unbiased reference, variance estimation is crucial for constructing confidence intervals, hypothesis testing, and comparisons across datasets. In this section, we derive a closed-form combinatorial formula for Var[*G*_*S*_(*r*)], eliminating the need for resampling over all 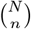 possible subsets. This derivation extends the expectation formula by incorporating second-order joint probabilities for pairs of points in *S*, allowing us to account for their co-occurrence in CSR estimation.

#### 3.1. Recap of the Expectation of *G*_*S*_(*r*)

As previously established, *G*_*S*_(*r*) is defined as:

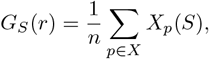

where *X*_*p*_(*S*) is an indicator variable that equals 1 if *p∈ S* and has at least one neighbor in *S* within radius *r*, and 0 otherwise. Using linearity of expectation, we obtained the exact expectation:

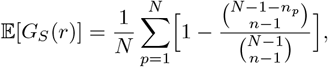

where *n*_*p*_ is the number of neighbors of *p* in *X*. This expectation captures the average nearest-neighbor behavior across all possible subsets of size *n*, but it does not describe the spread or variability of these values across different random subsets.

#### 3.2. Variance Definition and Reformulation

The variance of *G*_*S*_(*r*) is given by:

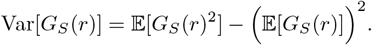

To compute 𝔼 [*G*_*S*_(*r*)^2^], we expand:

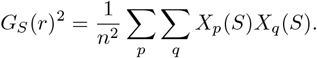

Taking expectations:

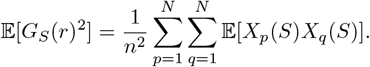

This sum consists of two terms:

1. Diagonal terms (*p* = *q*):

𝔼 [*X*_*p*_(*S*)*X*_*p*_(*S*)] = 𝔼 [*X*_*p*_(*S*)].

2. Off-diagonal terms (*p≠ q*):

𝔼 [*X*_*p*_(*S*)*X*_*q*_(*S*)] = Pr(*X*_*p*_(*S*) = 1 and *X*_*q*_(*S*) = 1), the probability that both *p* and *q* appear in *S* and have at least one neighbor in *S*.

Splitting these components:

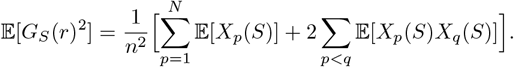

Subtracting (𝔼 [*G* (*r*)]) 2 gives Var[*G* (*r*)].

#### 3.3 Computing 𝔼 [*X*_*p*_(*S*)*X*_*q*_(*S*)] for *p≠ q*

To evaluate the off-diagonal terms, we introduce:

- *n*_*p*_: number of neighbors of *p*,
- *n*_*q*_: number of neighbors of *q*,
- *n*_*p,q*_: number of shared neighbors between *p* and *q*.

There are two cases:

**Case 1:** *p* **and** *q* **are direct neighbors (**dist(*p, q*) *< r*) If *p* and *q* are neighbors, then they automatically satisfy *X*_*p*_(*S*) = *X*_*q*_(*S*) = 1 whenever they are both in *S*, so:

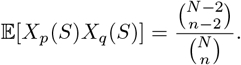

**Case 2:** *p* **and** *q* **are not direct neighbors (**dist(*p, q*) *≥r*) In this case, *p* and *q* must find other neighbors in *S*. The probability that neither finds a neighbor is:

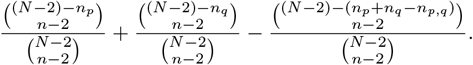

Thus,

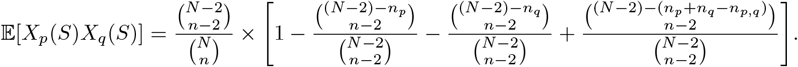

#### 3.4. Final Expression for Variance

Substituting into 𝔼 [*G*_*S*_(*r*)^2^]:

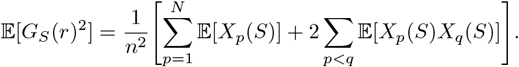

And the variance is:

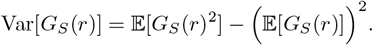

Alternatively, using covariance:

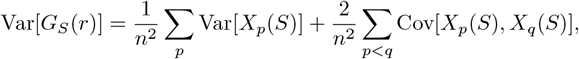

where:

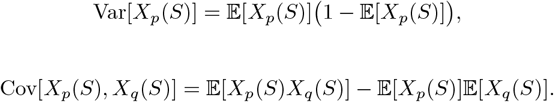

#### 3.5. Summary and Implications

This closed-form variance formula allows precise quantification of uncertainty in spatial clustering, enabling the construction of confidence intervals for CSR-corrected nearest-neighbor distributions. This eliminates the need for repeated permutations, making large-scale statistical comparisons of spatial clustering feasible. The ability to compute both the expectation and variance analytically strengthens the rigor of nearest-neighbor spatial analyses, making this approach invaluable in spatial omics and epidemiology. This derivation ensures that all combinatorial aspects of subset selection are handled exactly, bypassing the need to enumerate subsets while maintaining mathematical precision and computational efficiency. Importantly, however, is this solution scales poorly as the number of points increases, resulting in situations where permutations are more time efficient for identifying the null distribution under CSR (beyond ∼250 total points in a sample).

## Results

To evaluate the effectiveness of eliminating permutation-based estimation in nearest neighbor *G*(*r*) calculations, we applied our approach to a multiplex immunofluorescence (mIF) dataset of clear cell renal cell carcinoma (ccRCC). The dataset included a tissue microarray (TMA) containing a region of interest corresponding to a previously analyzed spatial transcriptomics sample. To control for spatial heterogeneity, we restricted the analysis to cytotoxic T cells (CD8+) in the tumor compartment, avoiding confounding effects from stromal regions (Figure 1).

**Figure 1:**
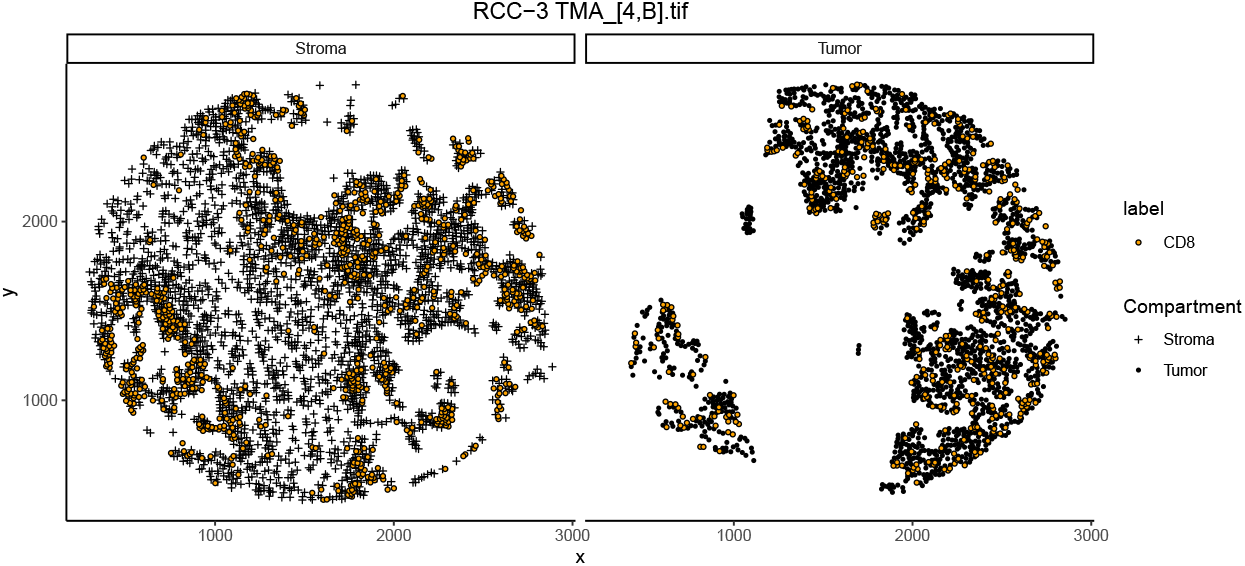
Example TMA core showing the spatial distribution of CD8+ cells in both the tumor and stroma compartments.

A key motivation for this work is the inaccuracy of theoretical CSR estimates in biological datasets where stationarity assumptions are violated. Theoretical CSR, which assumes a uniform spatial distribution of points within the window, overestimates clustering at small radii and underestimates clustering at larger radii [6], particularly in samples with uneven spatial structures such as tumors. To illustrate this, we computed both theoretical CSR and empirical CSR from 1,000 permutations. The results confirm that theoretical deviates substantially from empirical CSR when gaps and heterogeneities are present in the spatial distribution (Figure 2). This highlights the necessity of permutation-based CSR estimation to obtain an accurate null expectation against which to compare observed clustering patterns.

**Figure 2:**
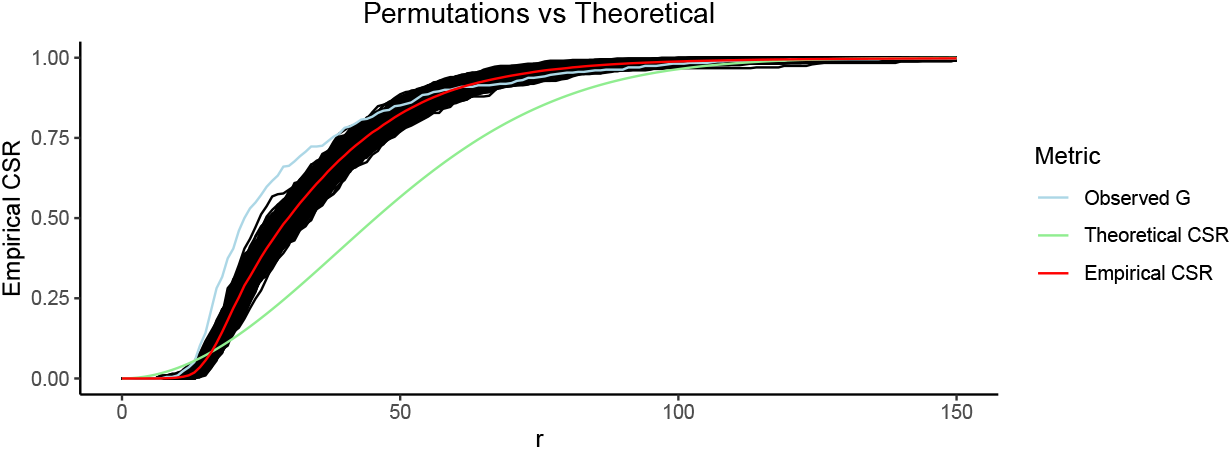
Theoretical *G*(*r*) overestimates clustering at low radii and under estimates clustering at larger radii in this sample (green line). *G*(*r*) calculated on 1000 permutations is shown as the black distribution of lines and mean of the permutations is shown in red. The *G*(*r*) of the true observed locations of cytotoxic T cells is shown as the light blue line, showing significantly greater clustering than expected (higher observed than the permutation distribution) between about *r* = 20 and *r* = 45. Axes are radii in pixels (x) and measured *G* across those radii (y).

To overcome the computational inefficiency of permutation-based approaches, we implemented an exact CSR calculation using a closed-form expectation that eliminates the need for iterative resampling. We compared this exact CSR to permutation-based CSR estimates across multiple levels of resampling, ranging from 100 to 10,000 permutations, and found that exact CSR is in line with the expected value obtained from permutations. This agreement demonstrates that permutations estimate CSR in varying degrees of accuracy, with 100 permutations the most inaccurate. Figure 3 shows different methods for calculating CSR (permutation and exact) with the CSR estimate of 100 permutations subtracted.

**Figure 3:**
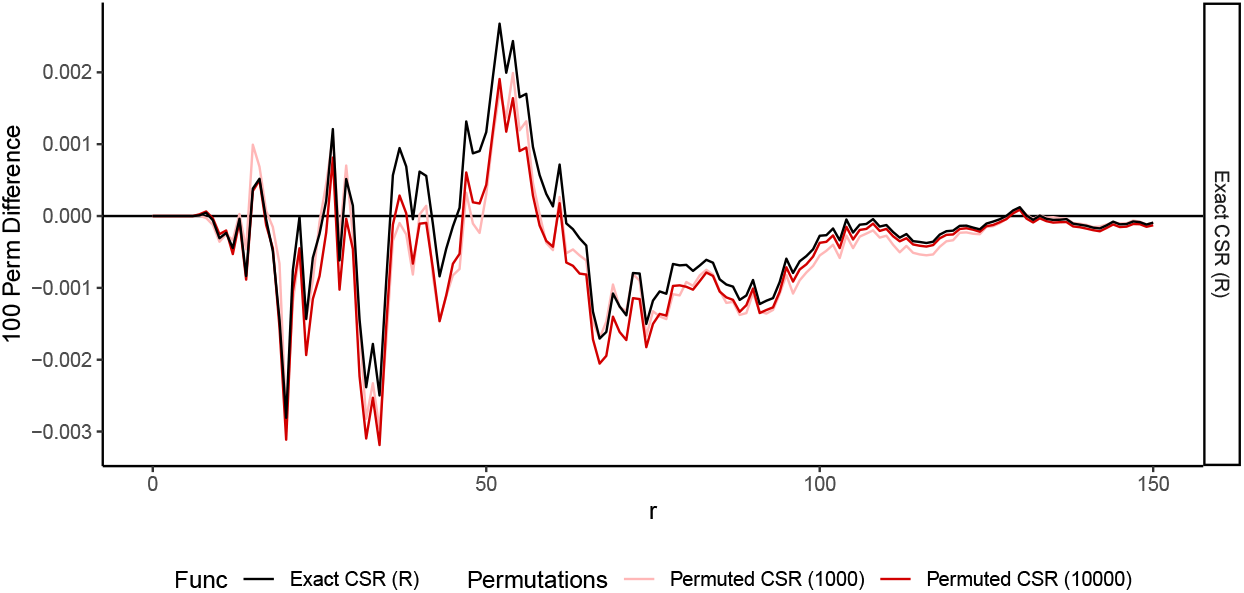
The difference of *G*(*r*) under complete spatial randomness compared to empirical estimate with 100 permutations. Permuted empirical estimates include mean of 1,000 permutations (pink) and 10,000 permutations (red), along with our exact approach (black) negating the need for permutations.

To further demonstrate the accuracy of our permutation-free approach, a sample of 30 points was simulated of which 5 were simulated ‘positive’ for a mark of interest (i.e., CD3+, CD19+, etc). All possible combinations of 5 points means that 142,506 measures of nearest neighbor *G*(*r*) need to be calculated and averaged in order to calculate the true CSR value for this sample. Figure 4 shows the result of averaging all combinations of 5 points (“Permuted CSR”) in this sample (red solid line). Calculating the exact CSR using our framework is shown in dashed purple, perfectly following the curve laid out by the averaging combinations. This shows that, without enumerating permutations, exact CSR is attainable both in decreased time needed to calculate as well as removing variation introduced through the permutation process.

**Figure 4:**
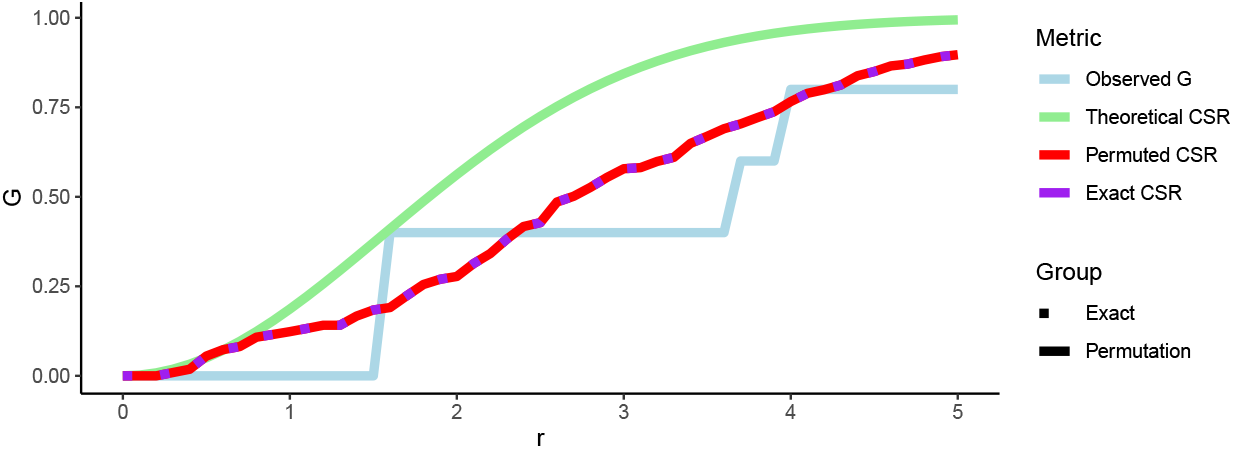
Exact complete spatial randomness compared to all combinations. A total of 30 points with 5 with a mark of interest were simulated followed by calculating the observed *G*(*r*) (light blue) as well as *G*(*r*) under theoretical CSR estimate (green), CSR calcualted from the mean of all combinations of 5 points (red solid line), and CSR using the exact method (purple dots; perfectly tracks the mean of all combinations of 5 points).

Beyond correctness, computational efficiency is a critical consideration for large-scale spatial analyses. We compared execution times and memory usage across multiple implementations of CSR estimation, including native R, R with Rcpp, and an optimized Rcpp implementation. As expected, permutation-based approaches exhibited poor scaling behavior, with computational cost increasing with the number of permutations. Again, as we saw in Figure 3, even 10,000 permutations for this sample deviates from the true sample-specific CSR. At higher permutation counts (e.g., 10,000), memory demands became prohibitive, limiting the feasibility of these methods for large data sets, especially when the number of points is greater than e.g. 10,000 (due to distance matrix size). In contrast, the exact CSR method reduced computational cost substantially, with the optimized implementation outperforming even permutation-based CSR with only 100 iterations. The most efficient version, implemented using an optimized Rcpp algorithm, provided a near-instantaneous solution with minimal memory overhead (mean time of 56.99ms vs 142.14ms for 100 permutations, and mean memory of 11.8MB vs 25.0MB for 100 permutations; Figure 5).

**Figure 5:**
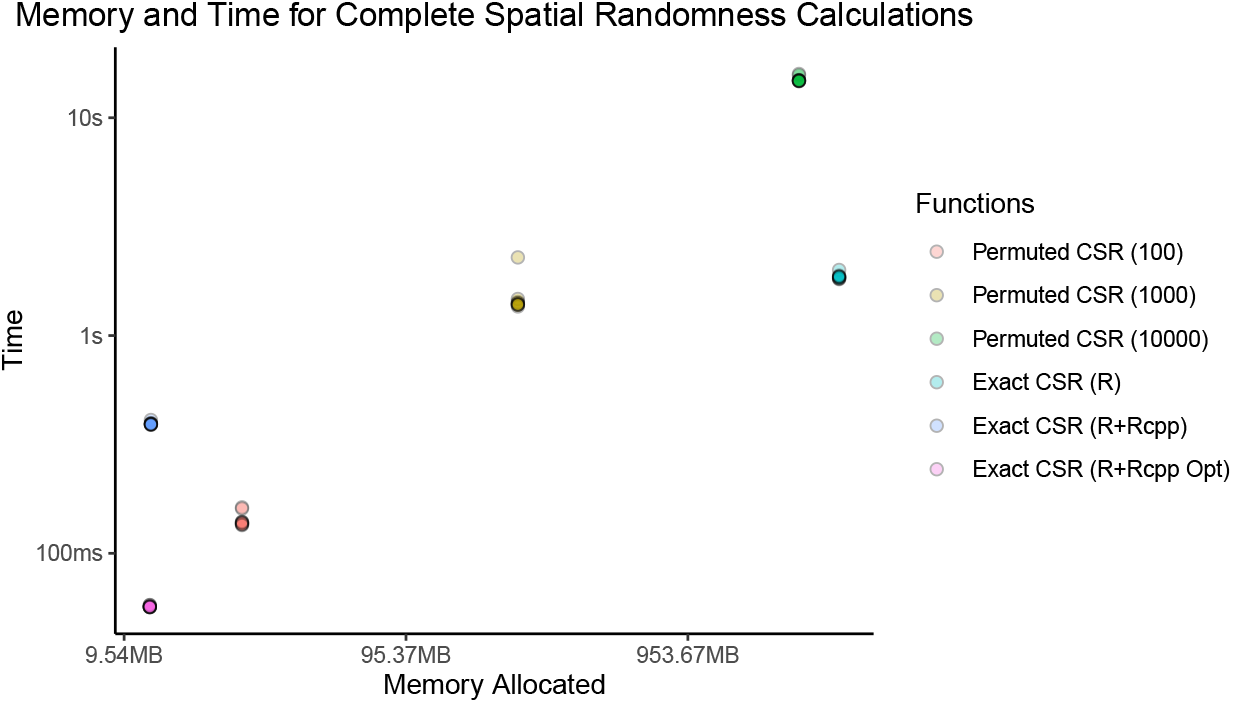
Memory usage and time to calculate the sample-specific *G*(*r*) of CSR.

These results confirm that exact CSR estimation offers a practical and scalable alternative to permutation-based methods. By eliminating the need for computationally expensive resampling while maintaining mathematical rigor, exact CSR enables efficient nearest neighbor *G*(*r*) calculations in complex spatial datasets. This improvement is particularly relevant for biological applications, where spatial stationarity assumptions are often violated, and permutation-based estimation can become a bottleneck. The optimized implementation ensures that large-scale spatial analyses can be performed both accurately and efficiently, facilitating more robust assessments of spatial clustering patterns in tissue microenvironments and beyond (Table 1).

**Table 1:** Memory usage and time to calculate empirical complete spatial randomness.

|  | Function | Time (mean) | Time (sd) | Memory (mean) | Memory (sd) |
| --- | --- | --- | --- | --- | --- |
| 1 | Permuted CSR (100) | 142.14ms | 1.03ms | 25.0 MB | 0 |
| 2 | Permuted CSR (1000) | 1.49s | 28.04ms | 237.8 MB | 0 |
| 3 | Permuted CSR (10000) | 15s | 41.68ms | 2.3 GB | 0 |
| 4 | Exact CSR (R) | 1.86s | 5.22ms | 3.2 GB | 0 |
| 5 | Exact CSR (R+Rcpp) | 393.88ms | 0.58ms | 11.9 MB | 0 |
| 6 | Exact CSR (R+Rcpp Opt) | 56.99ms | 0.05ms | 11.8 MB | 0 |

## Discussion

The exact derivation of the Nearest Neighbor *G*(*r*) function under complete spatial randomness (CSR) presented in this study represents an important methodological advancement in spatial point pattern analysis. By eliminating the need for computationally expensive permutation-based estimation, this approach provides a mathematically exact expectation of CSR while dramatically reducing computational overhead, both memory and speed. This improvement has uses across multiple scientific fields where spatial clustering and dispersion play a critical role especially when multiple samples that violate stationarity assumptions are being compared. Further, studies that use multiple samples where violation of the stationarity assumption may not be uniform, the theoretical estimation of CSR is inadequate for appropriate comparisons between samples. Traditional CSR estimation through permutations has been an essential but computationally costly step in spatial analyses, limiting its application to large datasets or studies requiring multiple subgroup comparisons. The ability to compute an exact expectation not only improves efficiency but also enhances reproducibility by eliminating the randomness associated with permutation-based approaches.

Nearest Neighbor *G*(*r*) has been widely used in disciplines such as ecology, epidemiology, and spatial omics, including multiplex immunofluorescence (mIF) and spatial transcriptomics [6,8–11]. With single-cell spatial omics, clustering is measured based on a subset of marks while we have locations for all cells in a sample. This provides a unique opportunity to use the true locations of cells in the sample rather than having to rely on where cells may lie under a poisson point process. Traditionally, CSR estimation relied on random permutations of observed spatial locations, requiring thousands of iterations to approximate an expected null distribution [6,10]. While effective, this approach is computationally prohibitive, particularly for large datasets or when multiple subgroup analyses are required. Our exact CSR derivation circumvents this limitation, yielding an expectation that aligns precisely with permutation-based means while executing orders of magnitude faster with significantly lower memory requirements. The computational burden of permutations often restricts the number of replicates researchers can feasibly run, leading to less precise estimations of CSR. The exact CSR method resolves this by providing a closed-form solution that is both mathematically rigorous and computationally efficient.

Our results demonstrate that theoretical CSR underestimates or overestimates clustering in heterogeneous tumor microenvironments, leading to misinterpretation of spatial organization. The necessity of sample-specific CSR estimation has been recognized in spatial statistics, but its implementation has been computationally expensive for large-scale studies. The identification of an exact solution provides a crucial improvement, ensuring that sample-specific spatial characteristics can be accounted for without excessive computational cost. Without this correction, observed clustering patterns may be exaggerated or suppressed depending on the spatial characteristics of the sample, leading to incorrect biological or epidemiological conclusions. By allowing precise CSR estimation, our method provides a necessary refinement that enhances statistical accuracy, reproducibility, and biological interpretability across spatial samples.

The optimized Rcpp-based implementation of the exact CSR method further enhances its practical utility. By leveraging efficient spatial indexing and numerical computations, this approach reduces computational time to near-instantaneous levels while requiring minimal memory overhead. This positions the exact CSR method as a scalable solution for high-throughput spatial analyses, allowing researchers to incorporate CSR corrections into routine workflows without compromising computational resources. In contrast to traditional methods that require researchers to balance computational feasibility with accuracy, this approach ensures that exact CSR values can be efficiently computed across many samples without sacrificing precision. As spatial datasets continue to increase in complexity and size, efficient methodologies like this will become increasingly essential for ensuring that statistical analyses remain feasible at scale.

### Limitations and Future Directions

While the exact CSR method provides a robust alternative to permutation-based approaches, several aspects warrant further exploration. First, the current formulation may exhibit scaling issues, especially for the variance. This means that in scenarios where it’s desired to identify significant deviations from CSR, only smaller samples (i.e., less than 250 points) can leverage this proposed framework effectively. For larger samples permuting positive cell or point labels is still more practical. Further optimizations with Rcpp or other languages, especially considering parallel processing, may provide drastic benefits. Second, this framework for the exact mean and variance of CSR considering a subset of points in a point pattern is applicable only in 2D space. With further developments in bioengineering, 3D tissue culture and imaging has been proposed but would not benefit from the current implementation. Lastly, this exact approach depends on cells or points of interest being a subset of a larger sample. Some ecological studies where all points belong to the same class (trees, rabbits, etc), this method does not apply and the theoretical CSR estimation is required.

## Conclusion

The development of an exact CSR formulation for nearest neighbor *G*(*r*) estimation represents a significant improvement in single-cell spatial point pattern analysis. By eliminating the need for permutations, this approach enhances both accuracy and computational efficiency, making CSR estimation feasible for high-throughput spatial datasets. The method’s applicability extends across multiple disciplines, offering a generalizable framework for assessing spatial organization. Given the growing reliance on spatial data in modern scientific research, ensuring that clustering assessments are both statistically rigorous and computationally feasible is essential. The exact CSR method achieves this by providing an analytically derived, reproducible, and scalable solution.

## Data Availability

Code to produce figures as well as functions (both in R and in Rcpp), are provided at https://github.com/ACSoupir/PermFree-NearestNeighborG. As part of our dedication to reproducible research and method sharing, I have create a wrapper function exact_g that extends spatstat::Gest to add the empirical CSR, variance associated with CSR, and p-value of the observed value deviating from CSR.

